# Glycoprotein K8.1 shapes KSHV liver and bone marrow tropism in the marmoset infection model

**DOI:** 10.1101/2024.10.31.621273

**Authors:** Christiane Stahl-Hennig, Berit Roshani, Anna K. Großkopf, Sarah Schlagowski, Samuel Alberto Mérida Ruíz, Ramona Vestweber, Eric Eymard, Charles Wood, Shanchuan Liu, Martina Bleyer, Nicole Stolte-Leeb, Stefan Pöhlmann, Denise Whitby, Armin Ensser, Alexander S. Hahn

**Author notes:** Current affiliation: HIV & AIDS Malignancy Branch, Center for Cancer Research, National Cancer Institute, Bethesda, Maryland, USA. Current affiliation: Labcorp, Kesselfeld 29, 48163 Münster (Samuel Alberto Mérida Ruíz). Current affiliation: University of Basel, Basel, Switzerland (Nicole Stolte-Leeb).

## Abstract

Kaposi’s sarcoma-associated herpesvirus (KSHV) is a human tumor virus. Common marmosets, a potential non-human primate model for vaccine development and infection studies, have previously been shown to support KSHV infection. We established infection of marmosets with BAC-derived, recombinant KSHV to analyze the role of individual glycoproteins (GPs) in tissue tropism. Three groups of four animals were infected intravenously with BAC-derived KSHV wild type (wt), EphA2-receptor-binding-deficient KSHV gH^ASAELAAN^, or KSHVΔK8.1, lacking glycoprotein K8.1. Viral DNA was detected in multiple tissues of all animals, with the highest loads in spleen, liver, heart, and bone marrow. In most tissues, viral DNA loads did not significantly differ between groups. However, KSHVΔK8.1-infected animals exhibited significantly lower KSHV DNA levels in liver and bone marrow. KSHV wt infection induced activation of CD4^+^ and CD8^+^ T-cells. In animals infected with KSHV wt, KSHV gH^ASAELAAN^ or KSHVΔK8.1, different B cell subpopulations expanded after infection, which suggests that the glycoproteins themselves or the cell tropism imparted by their presence or absence might shape the host response to KSHV. In summary, our findings demonstrate that neither the Eph receptor interaction nor GP K8.1 is essential for intravenous infection, but K8.1 critically contributes to KSHV tissue tropism in vivo.

## INTRODUCTION

KSHV was the last human herpesvirus to be discovered to date. Its nucleic acid was amplified from Kaposi’s sarcoma (KS) lesions and it is now well-established as the causative agent of KS.^1,2^ Both KSHV seroprevalence and KS incidence are comparatively low in most countries of the northern hemisphere, but are high in most parts of sub-Saharan Africa where KS ranks among the most common types of cancer.^2,3^ While most KS cases are currently associated with the HIV epidemic, KS had already been a well described entity in Africa before the advent of HIV. In particular, it was also known as a pediatric tumor, which is different from classic KS that occurs mostly in elderly individuals.^4^ KS also occurs in HIV-negative men who have sex with men at relatively younger ages.^5^ In addition to KS, the virus was later found to be associated with primary effusion lymphoma and a variant of multicentric Castleman’s disease, possibly also with osteosarcoma and with additional B cell entities, and it is associated with KSHV inflammatory cytokine syndrome (KICS), a severe inflammatory condition.^6–10^

In resource-limited regions, treatment of KS and other KSHV-associated diseases is challenging. Therefore, the development of a vaccine should be a priority. An animal infection model is crucial for such an endeavor. Firstly, it is needed for challenge studies. Secondly, it could be used to establish which glycoproteins or receptor interactions are critical for transmission and for establishing latent infection in specific target tissues such as the B cell compartment. Non-human primates can be excellent models for human diseases and are preferred models for vaccine development.

We therefore sought to build on a previous study by Chang et al.^11^ and establish the marmoset infection model for infection with recombinant BAC16-derived virus. The marmoset is an obvious choice for the development of a vaccine as it is evolutionarily closer to humans than the tree shrew, which was recently also demonstrated to support KSHV infection.^12^ The virus used by Chang et al. was rKSHV.219, a recombinant virus, based on the same KSHV strain as the commonly used BAC-derived KSHV BAC16.^13^ In contrast to rKSHV.219, BAC-derived virus is highly amenable to genetic manipulation and allows for expression of different transgenes which will be desirable for developing KSHV or its attenuated version as a viral vector. We made use of the full-length BAC16, carrying a BAC cassette that encodes a hygromycin resistance and GFP, to test whether KSHV encoding a large transgene can establish infection. Expression of larger transgenes and reporter genes can serve as a proof-of-principle for future applications, e.g. for the development of KSHV itself as a vaccine vector. We also sought to determine whether GFP expression would permit the monitoring of infection. Additionally, and most importantly, the use of the recombinant BAC16 system allowed us to investigate the impact of glycoprotein mutants on KSHV infection of marmosets.

How individual viral glycoproteins and their receptor interactions govern tissue tropism remains an open question for KSHV, with direct implications for which glycoproteins need to be included in a protective KSHV vaccine. This is true for herpesviruses in general, as they encode a complex set of glycoproteins. Glycoproteins B (gB), H (gH), and L (gL) together form the conserved herpesviral core fusion machinery.^14^ The gH/gL glycoprotein complexes of KSHV, of the related gamma-herpesvirus Epstein-Barr virus (EBV), and of the more distantly related human cytomegalovirus, a beta-herpesvirus, interact with EphA2 and other members of the Eph family of receptor tyrosine kinases to promote endocytosis, fusion, and infection.^15–22^ For KSHV the Eph interaction has been demonstrated to contribute to infection of a variety of cell types, as Eph family receptors are widely expressed, but it is not essential. KSHV or the related rhesus macaque rhadinovirus (RRV) remain infectious while bearing point mutations or deletions that abrogate the interactions with Ephs when applied at concentrations that are high enough to overcome the defect incurred by the missing Eph receptor interaction in vitro.^19,23^ RRV deleted in the locus coding for gL is even able to establish persistence in animals after experimental i.v. infection.^23^ Accessory glycoproteins like K8.1 of KSHV are not conserved between KSHV and more distantly related herpesviruses but still contribute significantly to infection, and compared to the engagement of Eph family receptors by gH/gL in a very cell-specific manner.^24–26^ Using this marmoset i.v. infection model, we compared two previously characterized virus mutants to wild type (wt) BAC16 virus with regard to host colonization. The first mutant, KSHV gH^ASAELAAN^, is unable to interact with the Eph family of receptors through mutation of a central interacting motif on gH and disruption of the association of gH with gL and exhibits reduced infectivity on Eph-expressing cells,^19,27^ and the second mutant, KSHVΔK8.1, does not express glycoprotein K8.1, which is a critical contributor to infection of certain B cells and keratinocytes, and exhibits reduced infectivity on these cells.^24,25,27^

## RESULTS

### Intravenous KSHV infection results in broad tissue distribution of viral DNA with highest viral genome load in liver, spleen, heart and bone marrow

After a lead-in period, which was used to establish baseline values, we inoculated four animals per group with 5.2×10^7^ infectious units of the respective, recombinant, BAC-<u>derived virus</u> in a volume of 500 µl (Fig. 1). Based on the comparison of intravenous and mucosal infection performed by Chang et al.,^11^ we opted for the intravenous route for inoculation to optimize achievable viral load. As the KSHV gH^ASAELAAN^ virus exhibits reduced infection *in vitro* on all analyzed cell types, likely due to the broad expression of EphA2 and related molecules, we decided to titrate the viruses on EphA2-positive SLK cells.^15,16,19,28^ Therefore, KSHV gH^ASAELAAN^ was used at higher virion concentrations compared to KSHV wt and KSHVΔK8.1 in order to compensate for its strongly reduced infectivity at the step of initial infection. *In vitro,* the defects of KSHVΔK8.1 are only apparent on very few cells and not on SLK.^24^ As the endpoint of our study was to identify tissue-specific contributions of the viral glycoproteins and their receptor interactions, we settled on this titration regimen to reach inoculation doses that assure sufficient take of the respective recombinant viruses. Animal 17358 in the wt group only received approximately one tenth of the inoculation dose subcutaneously as i.v. infection failed due to extremely thin saphenous veins. Animal 17312 received only 0.3 ml, animal 17430 received only 0.2 ml (both gH^ASAELAAN^ group), and animal 17365 (ΔK8.1 group) received only 0.3 ml of inoculum due to problems during i.v. injection.

**Fig. 1.**
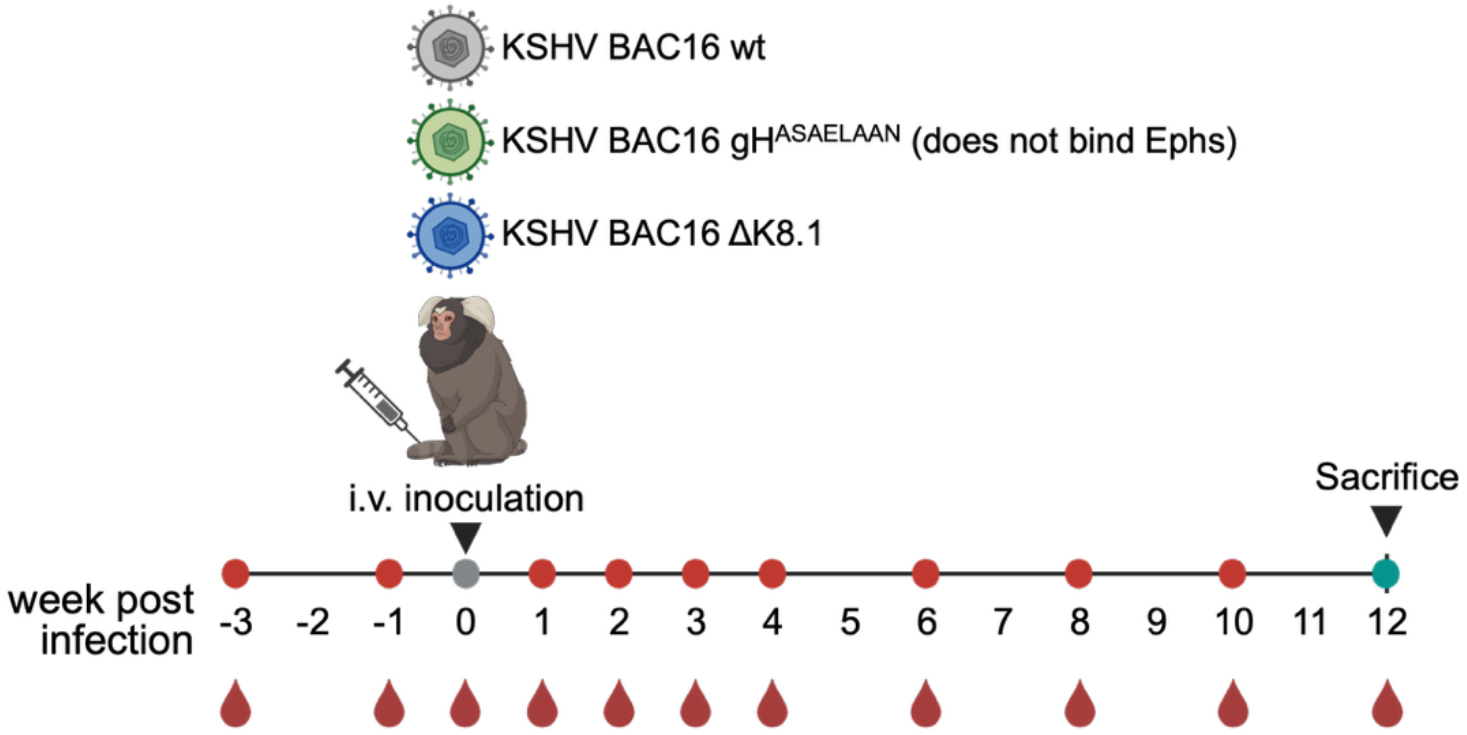
Experimental outline and schedule. Four common marmosets per group were infected with the indicated recombinant, BAC-derived viruses. This presentation was created in BioRender

The animals were followed for 12 weeks as outlined in Fig. 1. Virus detection from blood in general was unsuccessful except for single blips with most positive samples (eight) in the KSHV BAC16 gH^ASAELAAN^-infected group, compared to both other groups (two per group) (Fig. 2A). Therefore, at least within the timeframe of our sampling schedule, we did not detect consistent viremia. This does not exclude the possibility that a short viremic phase may have occurred between inoculation and e.g. day 7 or between day 7 and day 14 and was missed by our sampling schedule. The experiment was terminated at week 12 and the animals were sacrificed. Tissues were harvested and viral DNA load was determined.

**Fig. 2.**
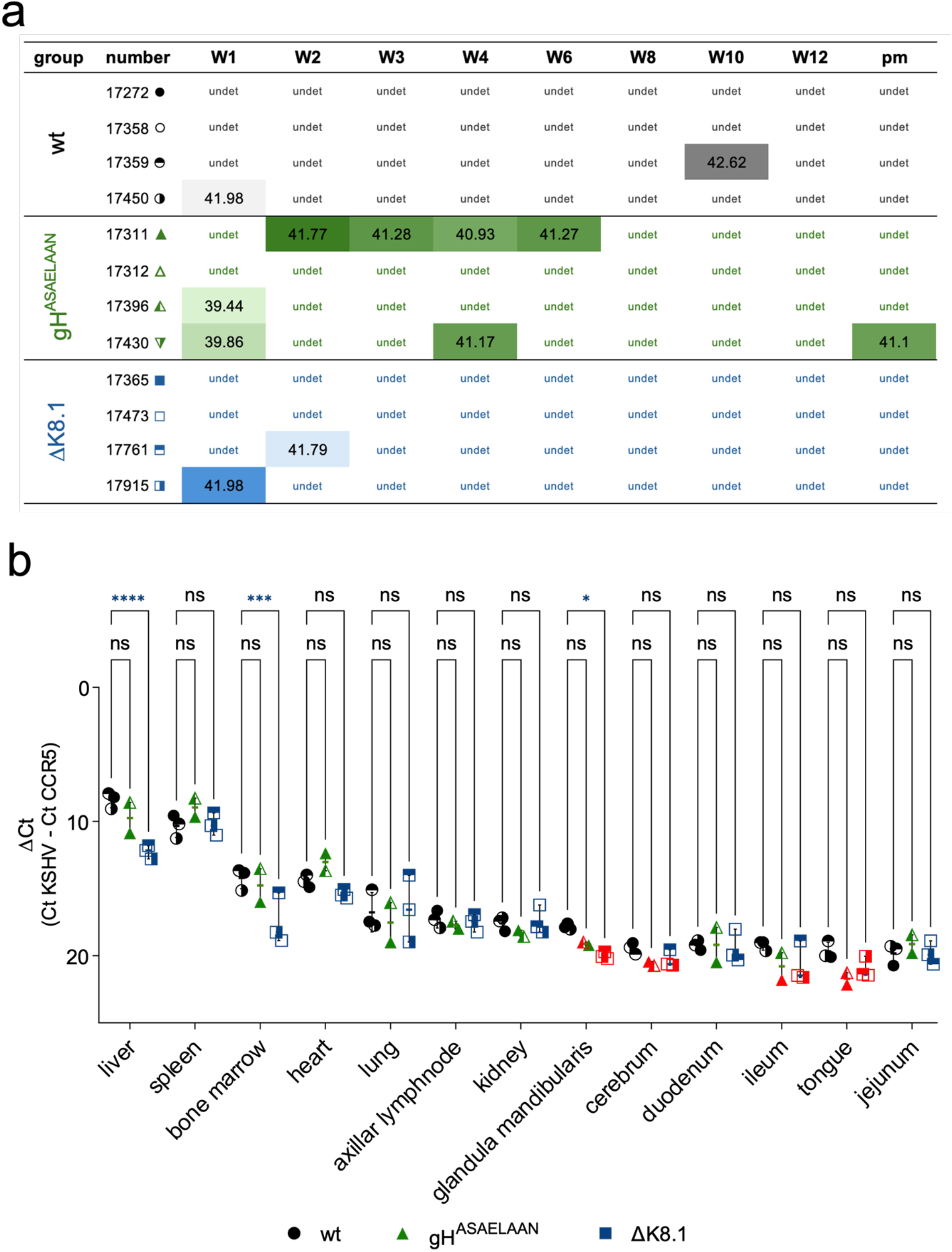
Viral DNA load in blood and various tissues. **a**, Blood was analyzed by qPCR at the indicated timepoints. Threshold cycle (Ct) values are shown. **b**, Tissues were harvested after sacrifice (approx. 12 weeks) and KSHV DNA and a cellular gene locus were quantified by qPCR. ΔCt values were calculated to quantify relative abundance; lower values indicate higher relative abundance. Each symbol represents one animal, red symbols indicate non-detects. Group means and standard deviations are shown. * p<0.05, *** p<0.001, **** p<0.0001, two-way ANOVA with Dunnett’s correction for multiple comparisons (wt n=3, gH^ASAELAAN^ n=2, ΔK8.1 n=3). W = week, pm = post mortem.

### KSHVΔK8.1 exhibits tissue-specific defects in liver and bone marrow

After sacrifice, tissues were harvested and viral DNA was quantified (Fig. 2b). For these analyses, animals that did not receive the exact amount of inoculum were excluded from the primary DNA tissue-load analysis as tissue viral loads may conceivably be sensitive to the amount of virus in the inoculum. Most tissues exhibited similar viral DNA loads after infection with KSHV wt, KSHV gH^ASAELAAN^ or KSHVΔK8.1. Noticeable exceptions were liver and bone marrow, in which KSHVΔK8.1 viral loads were significantly reduced when compared to KSHV wt. In the salivary gland, KSHV gH^ASAELAAN^ and KSHVΔK8.1 viral loads were visibly reduced compared to KSHV wt; this reached significance for KSHVΔK8.1 which was not detected in that tissue. In tongue samples KSHV gH^ASAELAAN^ and KSHVΔK8.1 viral DNA was not detected, and the difference in viral DNA load missed statistical significance for KSHV gHASAELAAN by a narrow margin (p=0.06). As viral DNA load was generally very low in these two tissues, it is unclear whether these findings are biologically meaningful. Assuming an average latent burden of ∼20 KSHV genome copies per cell, a compromise between reported numbers for KS and lymphoma cells,^29^ and near-optimal amplification efficiency approaching 2, we estimate an infection frequency on the order of approximately one infected cell per 10^4^ cells in tissues with the highest viral DNA loads. In case of occasional lytic replication with strong amplification of genome copy numbers, the frequency of infected cells would be even lower. In line with this estimate, we could not reliably detect LANA-positive cells by immunohistochemistry in liver tissue (not shown).

### KSHV infection of common marmosets results in seroconversion to the latency-associated nuclear antigen (LANA) of KSHV

We tested plasma of infected animals longitudinally for reactivity to LANA (Fig. 3). For these serological studies, we included all animals, irrespective of the actual amount of inoculum received. Three of the KSHV wt infected animals seroconverted, with peak values reached around week 3. One animal only exhibited weak seroreactivity to

**Fig. 3.**
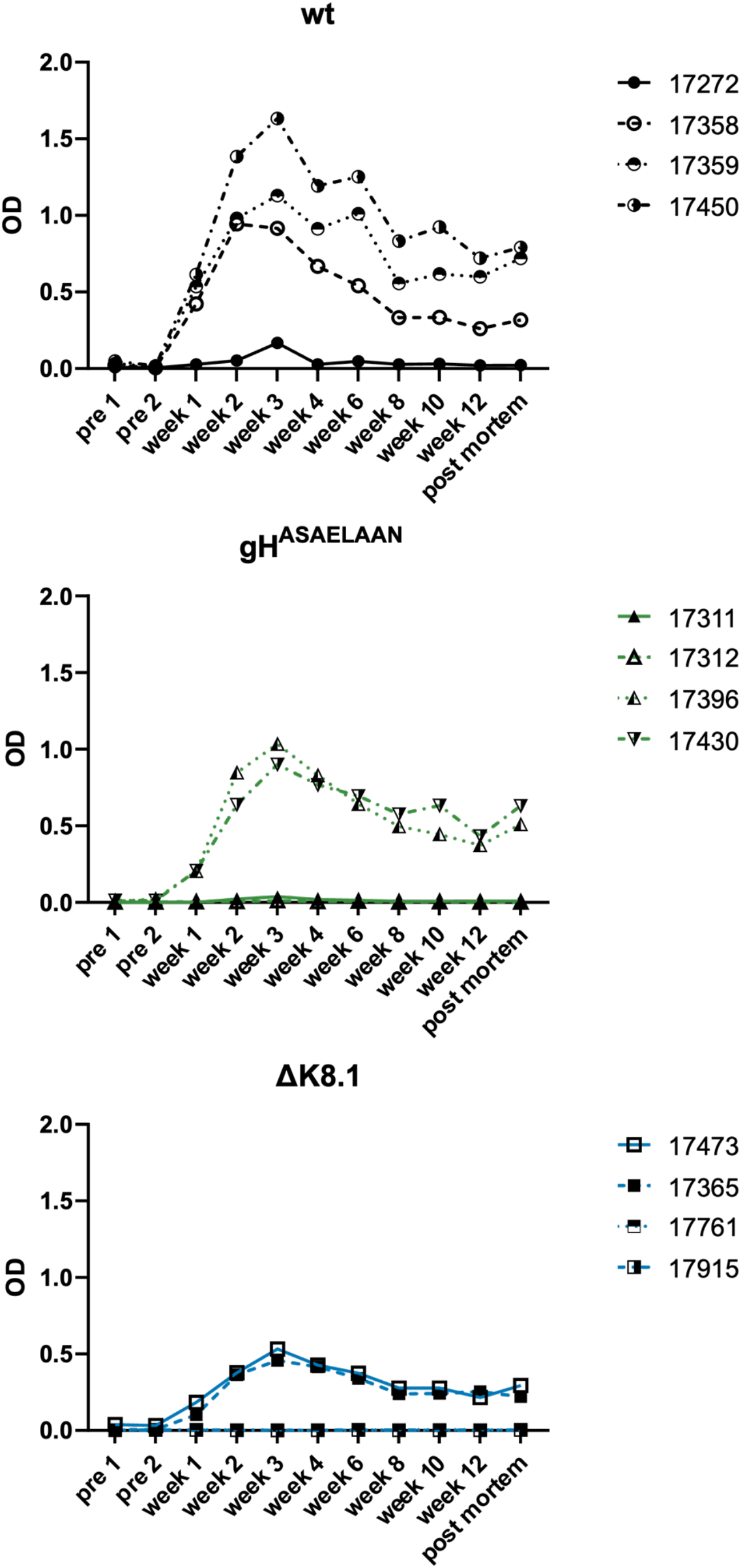
Seroconversion to the LANA antigen. Plasma obtained at the indicated timepoints was analyzed by ELISA for reactivity to KSHV LANA.

LANA, and only at the 3-week-timepoint after infection. In the KSHV gH^ASAELAAN^-infected group, two animals seroconverted to LANA with similar kinetics as observed for the KSHV wt group. In the ΔK8.1 group two animals seroconverted to LANA, and in these two animals the magnitude of the antibody response was lower compared to the other groups. However, given the high variance in responses within groups, this should be interpreted with caution. Overall, seroconversion to LANA did not obviously correlate with viral load.

### Infection of marmosets with KSHV results in the development of weakly neutralizing antibodies

We compared plasma samples taken before infection and at the 12-week-timepoint regarding their neutralizing activity. Since pre-infection plasma alone also exhibited an inhibitory effect at higher concentrations, we compared pre-infection and post-mortem plasma from each animal and calculated the relative infection at 1:500, 1:100, and 1:20 dilutions (Fig. 4a-c). All animals from the ΔK8.1 group but one developed at least weakly neutralizing activity (>20% neutralization at the lowest dilution). We next measured samples that had been collected across the whole 12-week period until sacrifice, which indicated that neutralizing activity developed rapidly in most animals. We found a considerable degree of variation between timepoints, which may be due to the small sample volumes combined with heat-inactivation and some precipitation occurring in the samples (Fig. 4d-f). To address this problem, we compared averaged results for the two pre-inoculation samples and for the five samples from week 6 to sacrifice for each animal (Fig. 4g). These results showed that only the KSHV wt-infected animals developed neutralizing responses that consistently inhibited by more than 50% at the 1:20 dilution tested and that were as a group different from the “no neutralization, relative infection = 1” null hypothesis with high significance. Neutralizing activity in animals from the two KSHV g^HASAELAAN^- and ΔK8.1-infected groups was more heterogeneous, and groupwise significance would not survive more stringent correction for multiple tests.

**Fig. 4.**
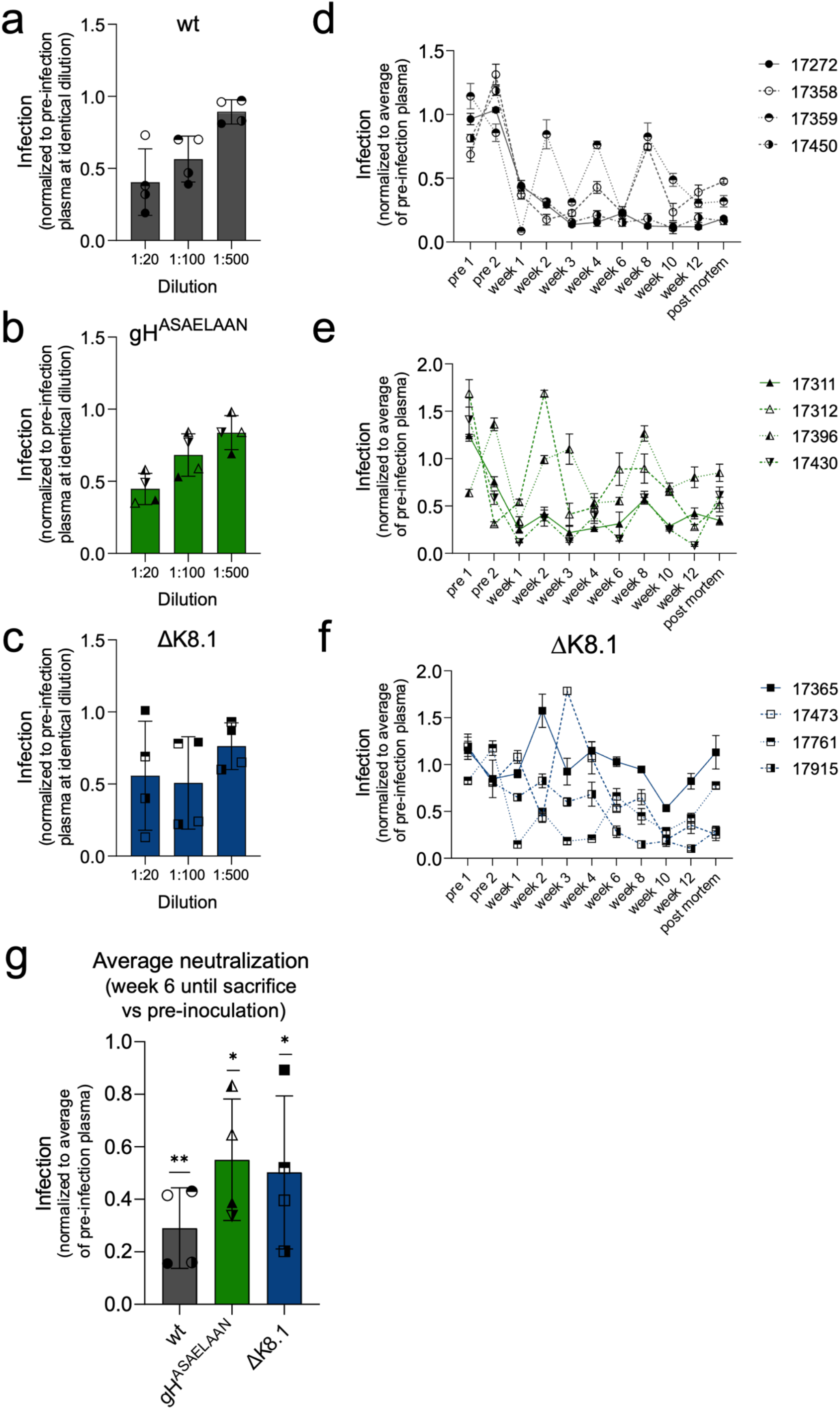
Development of neutralizing antibodies after experimental KSHV infection in marmosets. **a-c**, Pre-inoculation plasma or plasma obtained at sacrifice at the indicated concentration were mixed with KSHV BAC16 virus prior to infection of SLK cells. Infection was quantified and normalized to the pre-inoculation control. Group means and standard deviations within the group are shown. Each symbol represents the average of three replicate infections with plasma from one animal. **d-f**, Longitudinal plasma samples were analyzed as in **a** at 1:20 dilution. Error bars represent the standard deviation of three replicate infections per sample. **g**, For each animal, infection relative to the averaged pre-inoculation controls was averaged for all samples from week 6 to sacrifice, and group means were calculated. Means (n = 4 animals) and standard deviations are shown. One-sample t-test against 1 for each group, no correction for multiple comparisons * p<0.05, ** p<0.01.

We also performed a VirScan against linear epitopes from the KSHV proteome as described previously (Supplementary Fig. 2).^30^ It should be noted that these peptides are not representative of folded proteins. While increases in reactivity to peptides from some antigens, e.g., for ORF73 in animal 17450 were observed, we were not able to discern a consistent increase in reactivity towards certain KSHV-derived peptides.

### Common marmosets demonstrate signs of immune activation in response to KSHV infection

We analyzed the composition and activation status of the immune cell compartment of the infected animals over the time course of the experiment (Fig. 5). In the analysis, we included all animals. We were primarily interested in changes vs. baseline and the direction and magnitude of these changes, which are conceivably dictated by infection status and should be less sensitive to variations in the amount of the inoculum. In the BAC16 wt-infected animals, we observed significant activation of T cells as evidenced by an increase in PD-1-positive CD3^+^ cells as well as an increase in PD-1-positive CD8^+^- and PD-1-positive CD4^+^-T cells (Fig. 5a-c). A small variation of PD-1 positivity was also observed on CD20^+^ cells (Fig. 5d), but it did not reach significance. Interestingly, in the KSHV BAC16 gH^ASAELAAN^-infected animals and to a lesser degree in the KSHVΔK8.1 group, we observed a strong transient increase in CD21^+^/CD27^+^ CD20^+^ B cells (Fig. 5e), a population that was reported to represent memory cells that can proliferate at higher rates in rhesus macaques.^31^ While this increase did not reach significance for the KSHVΔK8.1 group, it was significant at four timepoints for the KSHV BAC16 gH^ASAELAAN^ group. We also observed an increase in CD20^+^/ CD21^-^CD27^-^ B cells compared to baseline in all groups (Fig. 5f), significantly at five timepoints in the KSHV BAC16 wt-infected animals, at three timepoints in the ΔK8.1 group, and at one timepoint in the g^HASAELAAN^ group. In humans, this particular B cell population has previously been associated with tissue-resident-memory B cells or a population of B cells that increase after prolonged pathogen exposure.^32^ There were a few instances where group means were temporarily but nevertheless significantly decreased in comparison to baseline, at single timepoints (Fig. 5 a, b, c, e). As these decreases were numerically small and not part of a larger trend, we did not further interpret them. As all T and B cell populations were GFP-negative, their increased frequency likely represented reactive expansion and not KSHV infection, which is also compatible with our inability to consistently detect viral DNA in blood. Overall, these findings demonstrate that experimental KSHV infection with BAC16 wt virus activated the animals’ immune systems over months.

**Fig. 5.**
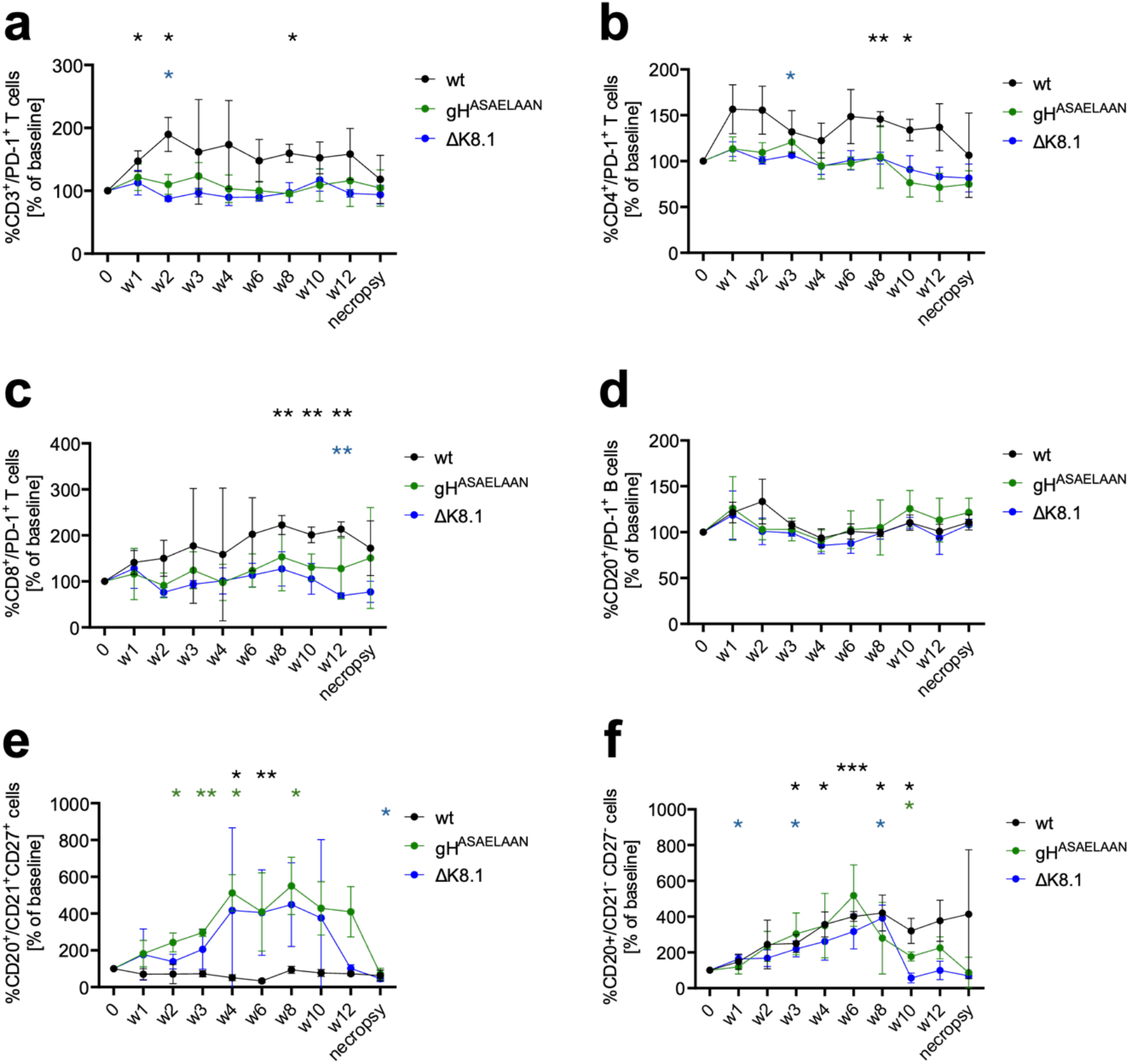
Immune cell populations after experimental infection with BAC16-derived KSHV wt, gH^ASAELAAN^ or ΔK8.1. For each group, values were compared to baseline. Timepoints that show a significant difference to baseline are indicated for each group by asterisks. **a**, PD-1 expressing CD3^+^ cells. **b**, PD-1 expressing CD4^+^ T cells. **c**, PD-1 expressing CD8^+^ T cells. **d**, PD-1 expressing CD20^+^ B cells. **e**, CD21/CD27 double-positive CD20^+^ B cells. **f**, CD21/CD27 double-negative CD20^+^ B cells. Significant changes to baseline for the KSHV wt, gH^ASAELAAN^ and ΔK8.1 groups are highlighted by asterisks. Group means and standard deviations are shown for each timepoint. Two-way ANOVA with Geisser-Greenhouse correction and Dunnett’s test, multiple comparison vs. baseline (week 0); * p<0.05, ** p<0.01, *** p<0.001.

## DISCUSSION

Our results demonstrate that BAC-derived KSHV establishes infection in common marmosets, which allows direct comparison of recombinant viruses generated in this reverse genetic system. Unlike in a previous study,^11^ infection was not accompanied by readily detectable viremia (Fig. 2a). Whether this reflects biological or technical variations is not entirely clear. We extracted DNA from whole blood samples, while Chang et al. analyzed purified PBMC. For PBMC-associated viral DNA, isolation of PBMC will lead to higher sensitivity, and the viral loads in PBMC reported by Chang et al. were close to the limit of detection, so they may have been slightly below our limit of detection.

Regarding the inoculation procedure, using a smaller volume in future studies may be beneficial as in several animals injection of the full inoculation dose of 500 µl proved problematic. Nevertheless, even after excluding some animals from the analysis, the tight grouping of viral DNA load in tissues allowed for meaningful analysis. The results obtained were not materially different from results obtained without censoring of animals with incomplete inoculation (Supplementary Fig. 1).

While the tropism of KSHV for spleen and bone marrow was expected for a lymphotropic herpesvirus, and a tropism for spleen is also in keeping with findings in the tree shrew model,^12^ the consistently observed high viral loads in liver and heart were unexpected. On the other hand, autopsy results of KS patients show that KS affects the liver in up to about one third of the cases,^33^ compatible with a certain tropism of the virus for that organ. KSHV tropism for the heart has not been described so far, but reports of cardiac involvement of KS exist.^34,35^ Our results suggest that further studies focusing on cardiac tropism of KSHV may be warranted. If the heart represents a target organ for KSHV in humans too, this may have consequences for infected individuals and should be addressed in future studies.

We observed significant activation of T cells in BAC16 wt-infected animals but not in animals infected with the two mutant viruses, which may indicate a certain degree of attenuation. While we found an increased frequency of CD20^+^/CD21^-^/CD27^-^ B cells in the animals of all groups (Fig. 5f), only KSHV BAC16 gH^ASAELAAN^-infected and KSHV BAC16 ΔK8.1-infected animals showed increased frequency of CD20^+^/ CD21^+^ CD27^+^ B cells, which was significant at four timepoints for the KSHV BAC16 gH^ASAELAAN^ group (Fig. 5e). This was rather unexpected as tissue viral loads in these animals were not significantly different from those in wt-infected animals, except for liver and bone marrow in the KSHVΔK8.1 group. Thus, interaction with cellular receptors, or the cell tropism dictated by these interactions, may shape the host response to KSHV infection. While this finding needs to be confirmed in future studies, it suggests that the interaction with Eph receptors, the only known receptors for the mutated region in gH/gL of the gH^ASAELAAN^ mutant, may exceed the described function in KSHV entry, as there were no obvious differences regarding the tissue tropism of the virus in our tissue panel but there apparently were differences in the cell populations that expanded reactively after infection.

Our finding that K8.1 is critical for liver and bone marrow infection highlights the importance of animal experiments. So far, *in vitro* experiments demonstrated that K8.1 is an important factor in the infection of some B cell populations and keratinocytes.^24,25^ Congruent with our results using K8.1 knockout virus, K8.1-targeting antibodies were effective at blocking infection of certain B cell lines *in vitro* in previous studies.^25^ Our finding that it contributes to bone marrow infection is in line with its role in B cell infection. K8.1’s role in infection of the liver, on the other hand, is unexpected, as was KSHV’s strong tropism for liver tissue, which should be explored further. Taken together, our findings identify K8.1 as a potential antigen of interest for vaccines to limit colonization of the liver, B cells, and bone marrow, even if infection cannot be fully prevented.

Importantly, our experimental design was intended to compare tissue colonization after successful inoculation rather than to quantify the contribution of Eph receptor binding to overall infectivity. Titration on EphA2-positive SLK cells therefore required a higher physical particle dose of the gH^ASAELAAN^ mutant. Our results consequently demonstrate that Eph receptor engagement is dispensable for tissue colonization following successful high-dose intravenous inoculation, but do not exclude an important role during natural transmission via the mucosal route or at limiting inoculum doses.

All animals exhibited relatively similar KSHV DNA viral load when comparing animals within one group or when comparing groups regarding viral DNA load in all tissues except for liver, bone marrow, and potentially also salivary gland and tongue.

Tongue was PCR-negative in both the KSHVΔK8.1 and gH^ASAELAAN^ group. Salivary gland was PCR-negative in the ΔK8.1 group. Both salivary gland and tongue were borderline positive in all animals from the wt group that had received the full inoculum. Viral DNA loads were highly similar within groups in one tissue and generally across most tissues, irrespective of seroconversion to LANA or neutralizing antibody response. This is particularly apparent in animals 17365 and 17396 from the KSHVΔK8.1 and gH^ASAELAAN^ group, respectively, which exhibited minimal neutralizing responses without discernible increase in viral loads (Supplementary Fig. 1). Animal 17365 showed in fact slightly lower viral loads and was one of the animals that had received slightly less inoculum, and while it developed low neutralizing responses, it exhibited seroconversion to LANA. VirScan revealed at most heterogeneous responses in a few animals, without a clear pattern (Supplementary Fig. 2), unlike what was observed in KS patients, where LANA protein harbors the most antigenic linear epitope.^30^ While only detecting reactivity to linear epitopes, this finding suggests that prolonged exposure to non-surface proteins like LANA or presentation of degraded viral antigens, which allows reactivity with linear epitopes, did not consistently occur in the animals that were tested.

Limited exposure to viral antigens would also be compatible with the variable degrees of seroconversion to LANA and is most compatible with rapid control of viral replication. The VirScan results may also generally reflect reduced breadth of the immune response after short-term infection over three months in the animals compared to long-term infection over years in the human host.

Overall, the absence of strongly neutralizing responses and the heterogeneity between animals in their responses suggest that T cell responses may be primarily responsible for the control of KSHV infection. This hypothesis is compatible with the known risk factors for KSHV-associated disease, in particular compromised T cell immunity as a consequence of HIV infection or immunosuppressive therapy in transplant recipients that targets T cells,^2^ and with the observation that antibodies elicited by natural infection are not consistently associated with protection against KS.^36,37^ This notion also fits well with our observation of PD-1 expression as a marker of T cell activation and potentially exhaustion, both of CD4^+^ and of CD8^+^ T cells (Fig. 5a-c). This finding has potential implications for a vaccine. While our results do not rule out that antibodies may protect against KSHV infection, and it seems intuitively plausible that high levels of neutralizing antibodies should afford some protection, our results together with the existing body of knowledge strongly suggest that a vaccine mediating non-sterilizing immunity should also induce T cell responses to limit host colonization by KSHV and prevent KSHV-associated pathogenesis, in addition to antibody responses against glycoproteins, including against K8.1.

## MATERIALS AND METHODS

### Animals, animal procedures, and animal housing

Three groups of four male common marmosets (*Callithrix jacchus*) were used. Two animals were housed together in one cage (2 pairs per group) according to the established animal welfare standards at the DPZ.

The infection protocol comprised inoculation of 500 µl concentrated cell culture supernatant containing 5.2×10^7^ infectious units by injection into the saphenous vein. Some animals could not be injected with the full dose when the vein would not support injection of the intended volume.

Infections and blood draws from the saphenous veins were performed under anesthesia. For anesthesia via injection, the animals were trained to retreat to their sleeping boxes from which they could be smoothly retrieved manually. An intramuscular injection was then administered comprising a mixture of 5% ketamine, 1% xylazine, and 0.1% atropine (GM-II, 0.1-0.15 ml/200 g body weight), providing sufficient depth of anesthesia and analgesia. The animals were subsequently placed in a sleep box to allow them to fall asleep.

For euthanasia, a mixture of 5% ketamine, 1% xylazine, and 0.1% atropine (GM-II) was administered intramuscularly (max. 0.1 ml/100 g body weight), providing sufficient depth of anesthesia and analgesia. Blood collection was then followed by an intraperitoneal administration of an overdose of pentobarbital (200-400 mg/kg body weight, equivalent to approximately 0.6 - 1.2 ml).

**Table 1.**
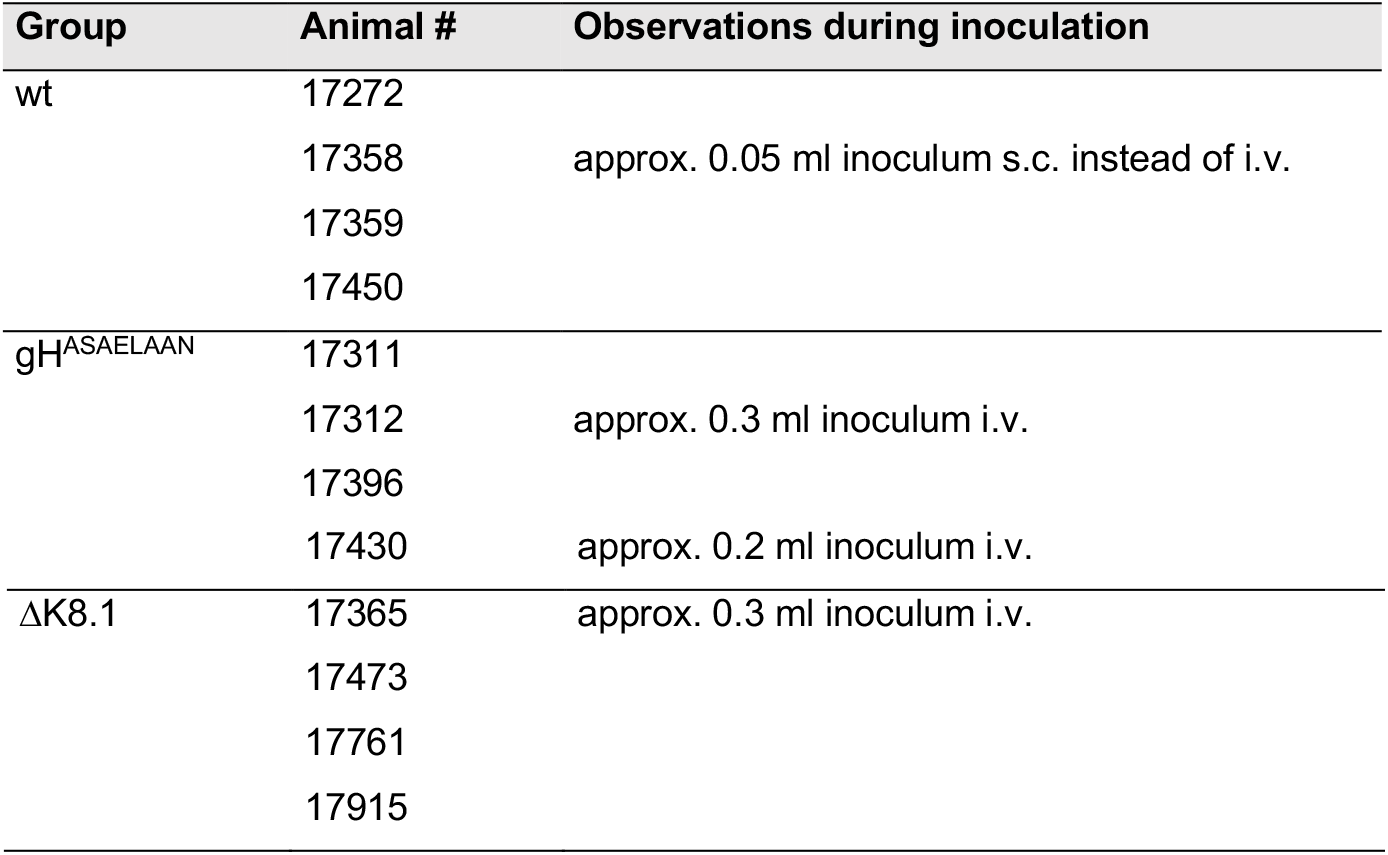
List of animals and deviations from experimental plan.

### Ethical approval statement

The study was approved by the relevant authorities (Niedersächsisches Landesamt für Verbraucherschutz und Lebensmittelsicherheit), license number 33.19-42502-04-20/3343.

The German Primate Center is permitted to house and breed non-human primates according to §11 of the German Animal Welfare Act. The permit was issued under license number 392001/7 by local veterinary authorities. Animals were housed under conditions in accordance with the German Animal Welfare Act, guidelines on the use of nonhuman primates for biomedical research by the European Union, and the recommendations of the Weatherall report.

### Viruses

Viruses were prepared as described previously from induced iSLK producer cells that had been transfected with the respective BAC constructs.^24,27^ The virus-containing supernatants were then concentrated by low-speed centrifugation in 50-ml-tubes overnight followed by careful aspiration of the supernatant down to the last approx. one milliliter. The virus was then resuspended, pooled and the procedure was repeated. Debris and precipitates were removed by brief high-speed centrifugation. The viruses were aliquoted and stored at -80°C. The inocula were titrated on SLK cells at different dilutions and the number of infectious particles was calculated using curve fitting and the Poisson distribution as described previously.^19^

### Neutralization assay

Plasma was heat-inactivated for 20 min at 56°C for experiments with heat-inactivation. The plasma was then preincubated with virus in cell culture medium at the respective dilutions for 30 min. For infection in 96-well plates, the cell culture medium was removed from the adherent SLK cells and the inoculum was added. Infection was quantified by measuring the percentage of GFP reporter gene positive cells as described before.^27^

### qPCR

Tissue DNA was isolated using the ISOLATE II Genomic DNA kit (Bioline) according to the manufacturer’s instructions using Proteinase K; DNA from blood was isolated using the DNeasy DNA Blood Mini kit (Qiagen) from 100 µl blood. KSHV BAC16 DNA was quantified using a primer/probe set directed against ORF59 and cellular DNA was quantified using a primer/probe set against CCR5 as described previously (KSHV ORF59 primer1 5’-GCCCACACCTTCCACTTCTAATA-3’, primer2 5’-GGTTAGCCTGGAGTCCTTAATC-3’, probe 5’-/56-FAM/CACTGTGTAA/ZEN/AGTCCCGGGTTGGTT/3IABkFQ/-3’).^19^ qPCR was performed on a StepOne Plus cycler (Thermo Fisher Scientific) in duplicate 20μl reactions using the SensiFAST Probe Hi-ROX Kit (Bioline) (cycling conditions: 3min initial denaturation at 95°C, 45 cycles 95°C for 10s and 60°C for 35s). Non-detects were set to Ct 42 as a conservative cut-off to strictly avoid over-interpretation of non-detects in the mutant virus groups, even if some single-positive duplicate PCR reactions were observed with Ct above 42. When only one duplicate well resulted in detection, that Ct value was used for the sample. Delta Ct values were calculated as delta Ct = (CtKSHV - CtCCR5). Where two samples were processed, delta Ct values were averaged.

### Serology

Plates were coated with recombinant LANA antigen as described previously and were stored at -80°C.^38^ The plates were thawed at room temperature (RT), washed three times with buffer (PBS-T, PBS with 0.02% Tween 20), and tapped dry.

Blocking was performed at 37°C for 30 min. After blocking, plates were washed three additional times. Subsequently, 50 µl of each sample, diluted in sample buffer (5% dry milk powder in PBS-T), was added to each well and incubated at 37°C for 2 hours. Plates were then washed three times and tapped dry, followed by the addition of 100 µl of conjugate diluted 1:12,000 in 2% dry milk powder in PBS (F(ab’)2 fragment goat anti-human IgG (H+L) peroxidase-conjugated secondary antibody, Jackson ImmunoResearch, article 109-036-088). This mixture was incubated at 37°C for 1 hour. After incubation, plates were washed five times and tapped dry before adding 100 µl of TMB substrate solution, which was incubated at RT for 10 min. The reaction was stopped with 1M H2SO4, and absorbance was measured at 450 nm with a reference wavelength of 630 nm.

KSHV serology results were further assessed by a phage immunoprecipitation assay (VirScan PhIP-Seq) as previously described.^30^ Here, post-mortem sera were used. Briefly, total IgG was quantified using monkey IgG ELISA (Immunology Consultants Laboratory #E-85G) according to the manufacturer’s instructions. The phage library was mixed with 2μg of IgG in duplicate and incubated prior to immunoprecipitation using Protein A and Protein G Dynabeads (Invitrogen, 10008D/9D). The bead mixture was then washed and remaining phage was lysed to release DNA contents. Finally, the DNA was indexed, gel-purified, and sent to the LSUHSC Translational Genomics core for quality check and sequencing. The sequencing was performed on an Illumina NextSeq2000 using the 100-cycle, single-end protocol. Average magnitude (i.e., the average frequency in which antibodies targeted the peptides spanning a viral protein, reported as minus log p-value or MLXP) against the KSHV proteome was calculated as shown in Supplementary Fig. 2. The MLXP values were also used to quality check the duplicate samples. Pearson correlation analysis of MLXP between replicates required a threshold of 0.7 to include marmoset samples in the final analysis. 8 of the 12 marmosets had both timepoints pass this quality check.

### Flow cytometry

The following monoclonal antibodies were used: CD3-AlexaFluor700 (clone SP34-2), CD4-V450 (clone L200) from BD Biosciences (Heidelberg, Germany), CD8-APC-Cy7 (clone HIT8a), CD27 – Brilliant Violet650 (clone O323), CD45-Biotin (clone 6C9), CD80 – PE-Dazzle (clone 2D10), HLA-DR-PerCP-Cy5.5 (clone L243), PD-1-APC (clone EH12.2H7) from BioLegend (San Diego, USA), CD20 – PE (clone H299), CD159a – APC (clone Z199.1) from Beckman Coulter (Krefeld, Germany) as well as CD21-SuperBright600 (clone HB5) from eBioscience (Frankfurt am Main, Germany). Antibodies were used at pretitrated concentrations. Prior to staining the anti-marmoset CD45-Biotin antibody was incubated with Streptavidin V500 (BD Biosciences, Heidelberg, Germany) for 15 min in the dark and then added to a mixture of the other monoclonal antibodies. 100μl of whole blood were stained with this mixture for 30 min at room temperature in the dark. Lysis of red blood cells (RBCs) and fixation were performed by incubating with RBC lysis/fixation solution (BioLegend, San Diego, CA) for 15 min.

Cells from the group infected with wild type virus were acquired using an LSRII cytometer (BD Biosciences). Cells from the other two groups were acquired using the ID7000 spectral analyzer from Sony Biotechnology. Analysis was performed using FlowJo 9.8.2 (Treestar, Ashland, OR) and GraphPad Prism Version 11.0.1. Comparable performance of the antibodies on the different flow cytometers was tested. Risk of confounding should be minimal as comparisons are made within each group and not between groups.

## AUTHOR CONTRIBUTIONS

C.S.H.: Designed and performed experiments, provided resources.

B.R.: Designed and performed experiments, analyzed data.

A.K.G.: Designed and performed experiments, analyzed data, edited the manuscript.

S.S.: Performed experiments.

S.A.M.: Performed experiments.

R.V.: Performed experiments.

E.E.: Performed experiments.

C.W.: Designed experiments and edited the manuscript.

S.L.: Validated data and edited the manuscript.

M.B.: Performed experiments.

N.S.L.: Designed experiments.

S.P.: Provided resources.

D.W.: Provided reagents.

A.E.: Validated reagents.

A.S.H.: Designed and conceived the study, analyzed data, wrote the manuscript.

## SUPPLEMENTARY INFORMATION

**Supplementary Fig. 1:**
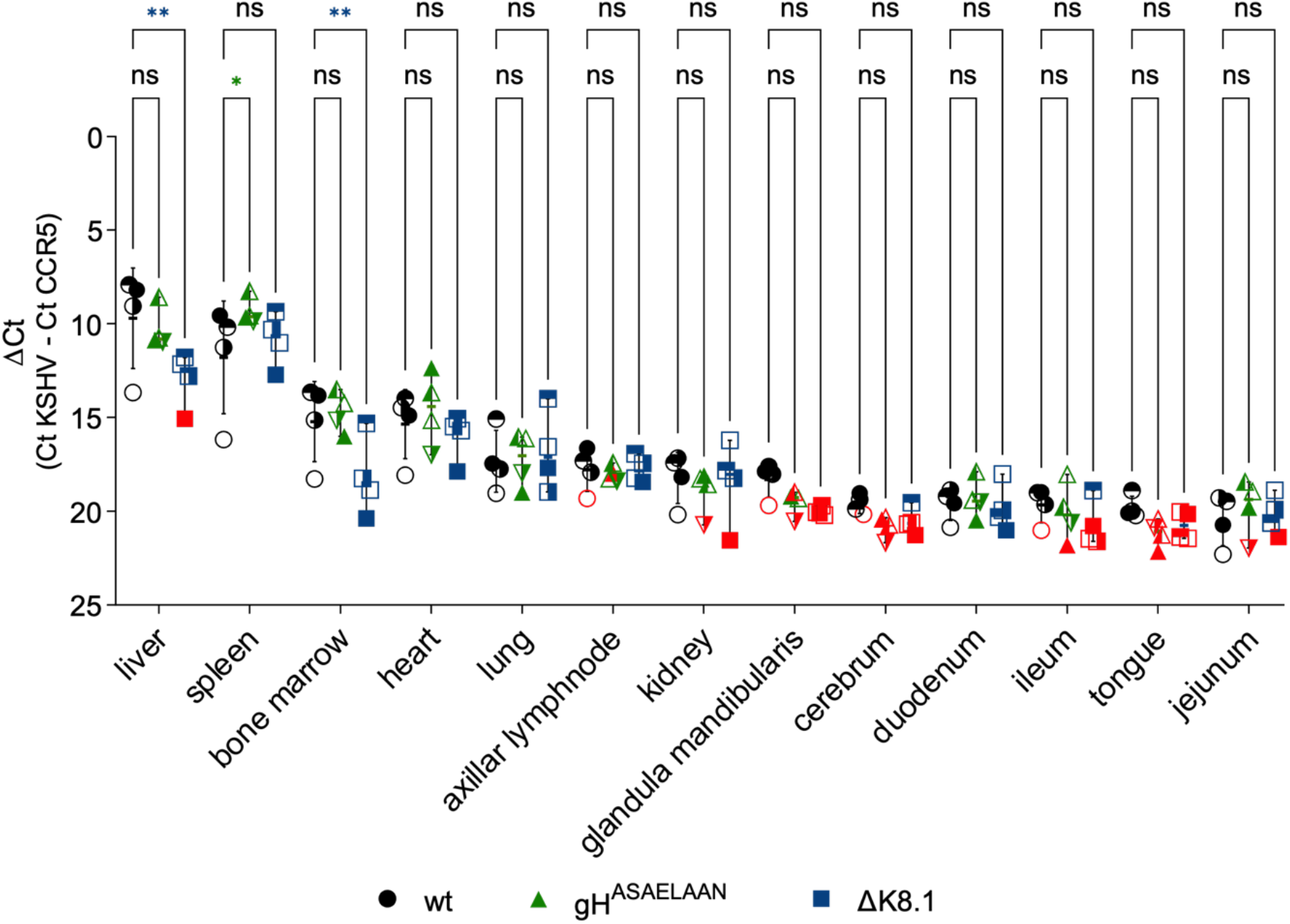
Viral tissue load analysis with all animals included. Tissues were harvested after sacrifice (approx. 12 weeks) and KSHV DNA and a cellular gene locus were quantified by qPCR. Delta-Ct values were calculated to quantify relative abundance. Red symbols indicate non-detects. Group means and standard deviations are shown. * p<0.05, ** p<0.01, two-way ANOVA with Dunnett’s correction for multiple comparisons (n=4 per group).

**Supplementary Fig. 2:**
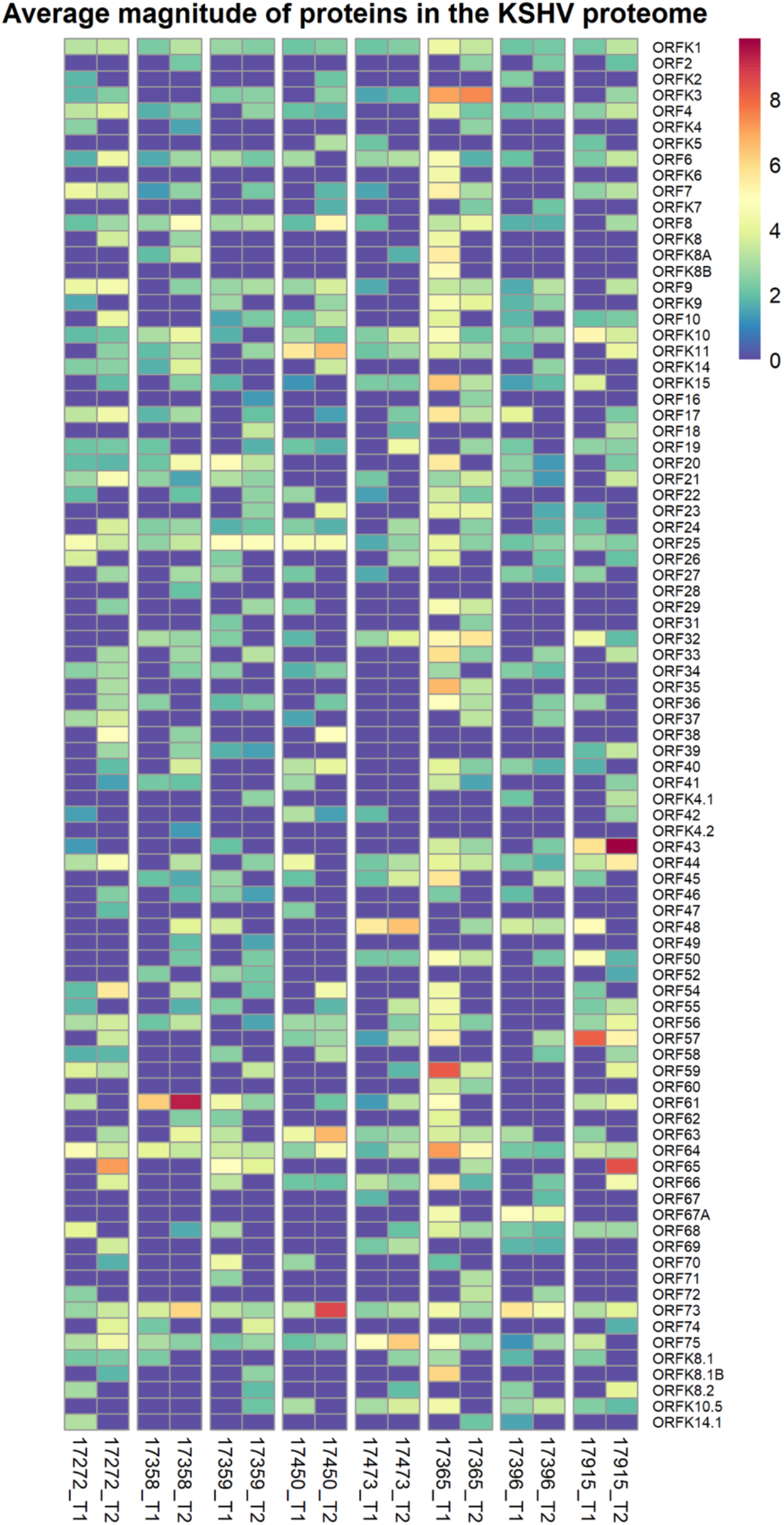
Peptide-Scanning analysis was performed with all KSHV open reading frames. Reactivity to the individual peptides is shown as heat map. T1 designates pre-infection plasma, T2 sera harvested at sacrifice (approx. 12 weeks). The scan was performed as described in Bennett SJ, Yalcin D, Privatt SR, Ngalamika O, Lidenge SJ, West JT, et al. Antibody profiling and predictive modeling discriminate between Kaposi sarcoma and asymptomatic KSHV infection. PLOS Pathogens. 2024;20: e1012023. doi:10.1371/journal.ppat.1012023.

